# Mapping the tRNA Modification Landscape of *Bartonella henselae* Houston I and *Bartonella quintana* Toulouse

**DOI:** 10.1101/2024.01.08.574729

**Authors:** Samia Quaiyum, Jingjing Sun, Virginie Marchand, Guangxin Sun, Colbie J. Reed, Yuri Motorin, Peter C. Dedon, Michael F. Minnick, Valérie de Crécy-Lagard

**Author notes:** Contributed equally (Samia Quaiyum, Jingjing Sun).

## Abstract

Transfer RNA (tRNA) modifications play a crucial role in maintaining translational fidelity and efficiency, and they may function as regulatory elements in stress response and virulence. Despite their pivotal roles, a comprehensive mapping of tRNA modifications and their associated synthesis genes is still limited, with a predominant focus on free-living bacteria. In this study, we employed a multidisciplinary approach, incorporating comparative genomics, mass spectrometry, and next-generation sequencing, to predict the set of tRNA modification genes responsible for tRNA maturation in two intracellular pathogens—*Bartonella henselae* Houston I and *Bartonella quintana* Toulouse, which are causative agents of cat-scratch disease and trench fever, respectively. This analysis presented challenges, particularly because of host RNA contamination, which served as a potential source of error. However, our approach predicted 26 genes responsible for synthesizing 23 distinct tRNA modifications in *B. henselae* and 22 genes associated with 23 modifications in *B. quintana*. Notably, akin to other intracellular and symbiotic bacteria, both *Bartonella* species have undergone substantial reductions in tRNA modification genes, mostly by simplifying the hypermodifications present at positions 34 and 37. *B. quintana* exhibited the additional loss of four modifications and these were linked to examples of gene decay, providing snapshots of reductive evolution.

## 1. Introduction

The conversion of genetic information from messenger RNA (mRNA) into functional proteins represents a fundamental and intricately regulated process in all living organisms^1^. At the heart of this complex mechanism, transfer RNA (tRNA) molecules function as indispensable adaptors, ensuring the precise and faithful translation of mRNA codons into amino acids^2^. Post-transcriptional modifications of tRNA molecules play pivotal roles in preserving the stability and integrity of tRNA molecules while enhancing the accuracy and efficiency of the decoding process^3^. As tRNA modification levels can be affected by stress or the metabolic status of cells, and these variations can affect the translation of specific mRNAs, specific modifications can be recruited to fulfill regulatory roles^4–7^. Even though such regulatory roles were postulated over 40 years ago^8^, it is only recently that the full regulatory or homeostatic loops have been dissected in different model organisms, including several pathogenic bacteria where they have been found to have roles in resistance to stress and virulence ^5,9–11^.

Exploration of the intricate regulatory mechanisms driven by tRNA modifications necessitates a comprehensive understanding of the entire set of modifications and the corresponding genes encoding the enzymes responsible for their installation. However, this knowledge remains largely incomplete, with significant gaps existing beyond a few well-studied model organisms, including the bacterial pathogen *Mycoplasma capricolum*^5^. Advancing our understanding requires the integration of analytical tools such as mass spectrometry (MS) or next-generation sequencing with comparative genomic analyses across diverse organisms spanning the tree of life^5^. Recent years have seen integrated studies that successfully cataloged tRNA modifications and associated genes in a few pathogenic bacteria, such as *Vibrio cholerae*^12^ and *Mycobacterium tuberculosis*^13^, as well as in the antibiotic-producing *Streptomyces albidoflavus*^14^. Notably absent from these investigations are members of obligate intracellular bacterial families like *Chlamydiaceae*, *Rickettsiaceae*, or *Ehrlichiaceae* which harbor significant pathogenic species for humans ^15,16^. Detecting tRNA modifications in intracellular pathogens presents unique challenges, including the low yield of tRNA during preparation and the potential for contamination by host cell tRNA. Yet, beyond unraveling the potential roles of tRNA modifications in the virulence of intracellular bacteria, mapping these modifications can contribute to our understanding of the genetic code’s evolution in organisms that undergo extensive genome reductions as part of their adaptations to specialized niches^17,18^.

*Bartonella henselae* Houston I, a member of the Hyphomicrobiales order, is a facultative intracellular Gram-negative pathogen that induces a spectrum of diseases, varying in severity. *B. henselae* is linked to ailments such as cat-scratch disease, which primarily affects children, and bacillary angiomatosis (BA), more commonly observed in HIV/AIDS patients^19,20^. Additionally, *Bartonella quintana* Toulouse, another facultative intracellular pathogen transmitted by human body lice, leads to trench fever and related illnesses^21,22^. These pathogens have complex life cycles as their reservoirs are mammalian vascular cells, but they are transmitted by arthropod vectors. *B. henselae* has evolved from an insect symbiont ancestor undergoing a first cycle of genome reduction^23^ with *B. quintana* undergoing a subsequent massive secondary genome reduction^24^. In addition, as these organisms can be cultured with vascular cells (erythrocytes, endothelial cells)^25^, they can be used as valuable test cases for employing the MS/NGS/comparative genomic approach to identify all genes related to tRNA modifications in intracellular bacteria.

Prior studies, which concentrated on rescuing the wobble base modification Queuosine (Q) in *B. henselae* and *B. quintana*, revealed that while the latter bacterium lost all genes associated with the pathway, *B. henselae* retained only two out of eight Q synthesis/salvage genes. These two genes constitute a basic salvage pathway, comprising a QPTR/YhhQ family transporter and a homolog of tRNA-guanosine (34) preQ_1_ transglycosylase (Tgt)^26^. Interestingly, this minimal pathway not only successfully salvages the Q precursor PreQ_1_ but also enables the rescue of the q base when present in high concentrations, a capability not shared by the *E. coli* orthologs. It is not clear which conditions would allow the salvage of these two precursors in natural environments, but analysis of tRNA extracted from *B. henselae* cells showed that preQ_1_ could be detected at the wobble position, a unique situation, to date. The current study focuses on predicting the remaining tRNA modifications to generate a better picture of decoding in these organisms.

## 2. Methods

### 2. 1. Bioinformatic Analyses

The genomic sequences of *Bartonella henselae* str. Houston I ((NCBI Taxon ID 38323; NC_005956.1) and *Bartonella quintana* str. Toulouse ((NCBI Taxon ID 283165; NC_005955) were used for all analyses. tRNA isoforms information was obtained from GtRNA database^27^ (http://gtrnadb.ucsc.edu/GtRNAdb2/genomes/bacteria/Bart_hens_Houston_1/ and http://gtrnadb.ucsc.edu/GtRNAdb2/genomes/bacteria/Bart_quin_Toulouse/). The BV-BRC platform ^28^ was used for protein family identifications, physical clustering analyses, and species tree construction using FastTree (version 47). Species trees were also generated using PhyloT (https://phylot.biobyte.de) (database version 2023.2) and Protein families were also identified with KO numbers in the KEGG database^29^. UniProt was used for ID mapping and advanced search tools^30^. NCBI was used for BlastP analyses^31^ and literature searches^32^. GizmoGene (http://www.gizmogene.com/) was used to make the physical clustering figures and Modomics^33^ for analyses of tRNA sequences. iToL (https://itol.embl.de) (version 6.8.1) ^34^ and MORPHEUS (https://software.broadinstitute.org/morpheus) were used to visualize the presence/absence data of protein families along the species trees.

### 2.2 Strains, Media and Bulk tRNA Preparation

*B. henselae* Houston I was obtained from the American Type Culture Collection (ATCC 49882). *B. quintana* Toulouse was a generous gift from Volkhard Kempf (Goethe-Universität Frankfurt). Both species were cultivated as previously described^25^on HIBB agar plates [i.e., Bacto heart infusion agar (Becton, Dickinson, Sparks, MD) supplemented with 4% defibrinated sheep blood and 2% sheep serum (Quad Five, Ryegate, MT) by volume] for 4 days (*B. henselae*) or 10 days (*B. quintana*) at 37°C, 5% CO_2_ and 100% relative humidity. Following harvest into ice-cold heart infusion broth, tRNAs were collected from the bacterial cells as previously described^26^.

### 2.3 LC-MS Analysis of Nucleosides in tRNA Samples

tRNA for each sample [3 µg (S1-S3 in Table S4a) or 1.8 µg (S1-S5 in Table S4b)] was hydrolyzed in a 50 µL or 30 µL digestion cocktail containing 12.5 U or 2.49 U benzonase, 5 U or 3 U CIAP (calf intestinal alkaline phosphatase), 0.15 U or 0.07 U PDE I (phosphodiesterase I), 0.1 mM deferoxamine, 0.1 mM BHT (butylated hydroxytoluene), 5 ng or 3 ng coformycin, 50 nM or 25 nM ^15^N-dA (internal standard [^15^N]_5_-deoxyadenosine), 2.5 mM MgCl2 and 5 mM Tris-HCL buffer pH 8.0. The digestion mixture was incubated at 37 °C for 6 h. After digestion, all samples were analyzed by chromatography-coupled triple-quadrupole mass spectrometry (LC-MS/MS). For each sample, 600 ng of hydrolysate was injected for each of the two technical replicates. Using synthetic standards, HPLC retention times of RNA modifications were confirmed on a Waters Acuity BEH C18 column (50 × 2.1 mm inner diameter, 1.7 µm particle size) coupled to an Agilent 1290 HPLC system and an Agilent 6495 triple-quad mass spectrometer (**Table S1**). The Agilent sample vial insert was used. The HPLC system was operated at 25 °C and a flow rate of 0.35 mL/min or 0.3 mL/min in a gradient (**Table S2** and **Table S3** with Buffer A (0.02% formic acid in water) and Buffer B (0.02% formic acid in 70% acetonitrile). The HPLC column was coupled to the mass spectrometer with an electrospray ionization source in positive mode with the following parameters: dry gas temperature, 200 °C; gas flow, 11 L/min; nebulizer, 20 psi; sheath gas temperature, 300 °C; sheath gas flow, 12 L/min; capillary voltage, 3000 V; nozzle voltage, 0 V. Multiple reaction monitoring (MRM) mode was used for detection of product ions derived from the precursor ions for all the RNA modifications with instrument parameters including the optimized collision energy (CE) optimized for maximal sensitivity for the modification. Signal intensities for each ribonucleoside were normalized by dividing by the sum of the UV signal intensities of the four canonical ribonucleosides recorded with an in-line UV spectrophotometer at 260 nm.

### 2.4 tRNA Modification Analysis by Next Generation Sequencing

Analysis of tRNA modifications present in *B. henselae* tRNA fractions was performed by a combination of three previously published original protocols, namely RiboMethSeq, AlkAnilineSeq and HydraPsiSeq. Additional analysis of eventual m^5^C residues was done by standard RNA bisulfite sequencing.

#### 2.4.1 RiboMethSeq

RiboMethSeq protocol allowed us to map 2′-O-methylations by assessing their protection against alkaline cleavage^35,36^. To further enhance our analysis, we also extracted reverse transcriptase (RT) misincorporation signatures at RT-arresting nucleotides^37,38^. As part of the RiboMethSeq tRNA analysis, total RNA from *B. henselae* was fragmented under alkaline conditions, followed by library preparation and sequencing^35^. The RT misincorporation signatures were derived from the Samtools mpileup format and were manually verified by examining the aligned reads in the *.bam file using the Integrated Genome Viewer (IGV)^39^.

#### 2.4.2 AlkAnilineSeq

Analysis of m^7^G and D modifications in *B. henselae* tRNAs was done by AlkAnilineSeq protocol^40,41^. This method also detects m^3^C and ho^5^C, but these residues have not been reported in bacterial tRNAs. Eventual cleavage signals may be also observed for s^2^C and ho^5^U intermediates at position 34 of tRNAs. This method exploits sensitivity of certain RNA modified bases to ring-opening at high temperature and alkaline conditions, the resulting damaged base or RNA a basic site is further cleaved by aniline and adapter is ligated to the released 5’-P termini.

#### 2.4.3 HydraPsiSeq

For the mapping of pseudouridine (Psi) modifications, we employed a chemical-based protocol using hydrazine cleavage, termed HydraPsiSeq^42–44^. This protocol measures resistance of pseudouridines and m^5^U (rT) residues to the hydrazine cleavage. Subsequent aniline treatments convert damaged U residues to cleavages of the polynucleotide chain. Ligation of the sequencing adapter is done as in the AlkAnilineSeq protocol (see above). Of note, k^2^C (lysidine) present in bacterial tRNA^Ile^_CAU_ at the wobble position shows extensive cleavage upon hydrazine treatment and thus can be detected as positive cleavage signal.

#### 2.4.4 RNA bisulfite sequencing (RBS)

Analysis of eventual m5C modifications in *B. henselae* tRNAs was performed with the well-established bisulfite conversion protocol developed by Schaefer et al^45^. The EZ RNA Methylation Kit (Zymo Research #R5001) was used for RNA bisulfite treatment followed by desulphonation. Bisulfite-converted RNAs were then end-repaired and subjected to NEBNExt Small RNA Library Prep Set for Illumina (NEB, E7330L), according to manufacturer’s recommendations. Sequencing was performed in SR50 mode, with sequencing reads aligned to C->T converted reference sequence for *B. henselae* tRNAs. 3.1

## 3. Results and Discussion

### 3. 1. Gathering tRNA Gene Sets And tRNA Modification Nucleoside Profiles for Two *Bartonella* Species

The *B. henselae* genome encodes a total of 43 predicted tRNA genes, specifying 38 distinct iso-acceptors. In comparison, the genome of *B. quintana* comprises 42 tRNA genes with an equivalent number of iso-acceptors, albeit with the loss of one of the triplicate copies of initiator-tRNA^Met^_CAU_ (**Fig. 1**). This reduction in iso-acceptors is modest when compared to *E. coli*, with only four losses: tRNA^Arg^, tRNA^Gly^, tRNA^Pro^, and tRNA^SelCys^. This contrasts sharply with organisms with reduced genomes, such as mollicutes, which can pare down the number of tRNAs to 28^46^. Notably, while the total number of tRNA genes in *E. coli* is 86, often present in five copies ^47^, the two *Bartonella* species have significantly reduced the duplicate gene copies of any given iso-acceptor (**Fig. 1**).

**Figure 1.**
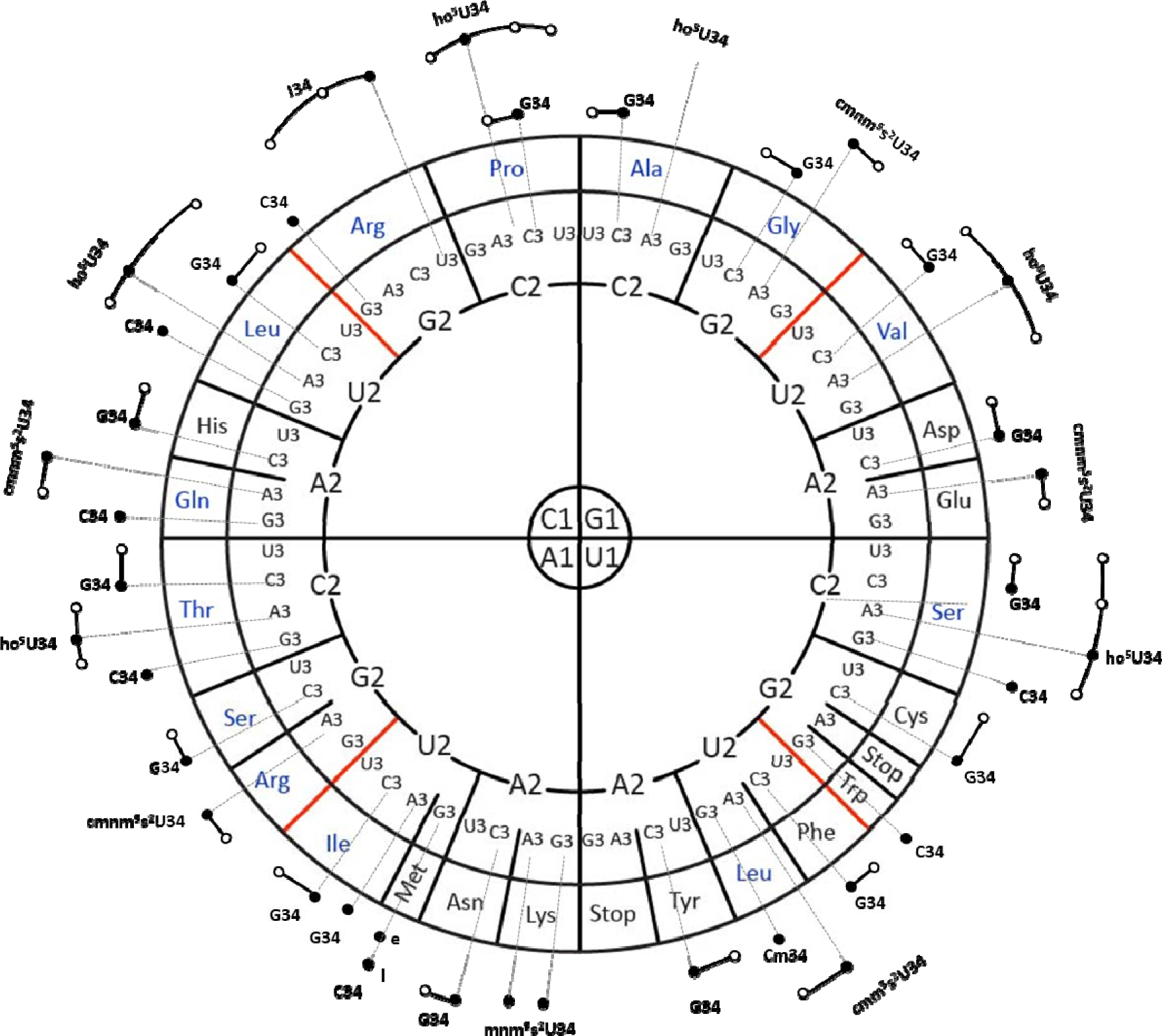
tRNA isoacceptors genes; a distribution for *Bartonella henselae* Houston I. A circular representation of the genetic code, highlighting modified nucleosides, is presented, following the format described by ^50^. In this representation, each amino acid codon sequence i read from the inside out (1-2-3). The dark dots represent the tRNA isoforms encoded by the genome. Additionally, the elongator or initiator tRNA^Met^_CAU_ are denoted by “e” and “i,” respectively. The predicted modifications of the ASL of any given isoacceptor in *B. henselae* and the predicted expanded decoding capacity through wobble or expanded wobble of each isoacceptor are noted by dotted lines and white circles.

In the absence of post-transcriptional modifications, these sets of tRNAs would be incapable of decoding the 64 codons^48^. To illustrate, tRNA^Ile^_CAU_ requires the modification of cytosine to lysidine to decode AUA codons. However, the modification status of tRNA molecules from *Bartonellaceae* was unknown, as none had been previously sequenced^33^. Consequently, we undertook an LC-MS/MS analysis, examining the ribonucleosides obtained from the enzymatic digestion of bulk tRNA extracted from both *Bartonella* species cultivated on HIBB plates, as detailed in the methods section. Six independent tRNA samples from *B. henselae* and two from the more challenging-to-culture *B. quintana* were analyzed, leading to the detection of 31 modified ribonucleosides in these samples, 30 with standards and 1 (ms^2^t^6^A) without but with genetic evidence as discussed in the section below (**Table 1** and **Table S4a-b**). However, not all modified ribonucleosides in this list are derived from *Bartonella* tRNAs. Indeed, rRNA modifications are routinely detected in analyses of tRNA samples as we previously discussed when analyzing tRNA modification profiles in *Bacillus subtilis*^49^ and in the current analysis we can also detect modifications both from rRNA or tRNA from the sheep cells present in the culture media. To predict the tRNA modification profiles of the *Bartonella* species specifically, we combined the results of LC-MS with bioinformatic analyses and NGS-based modification detection methods as described in the next sections.

**Table 1:**
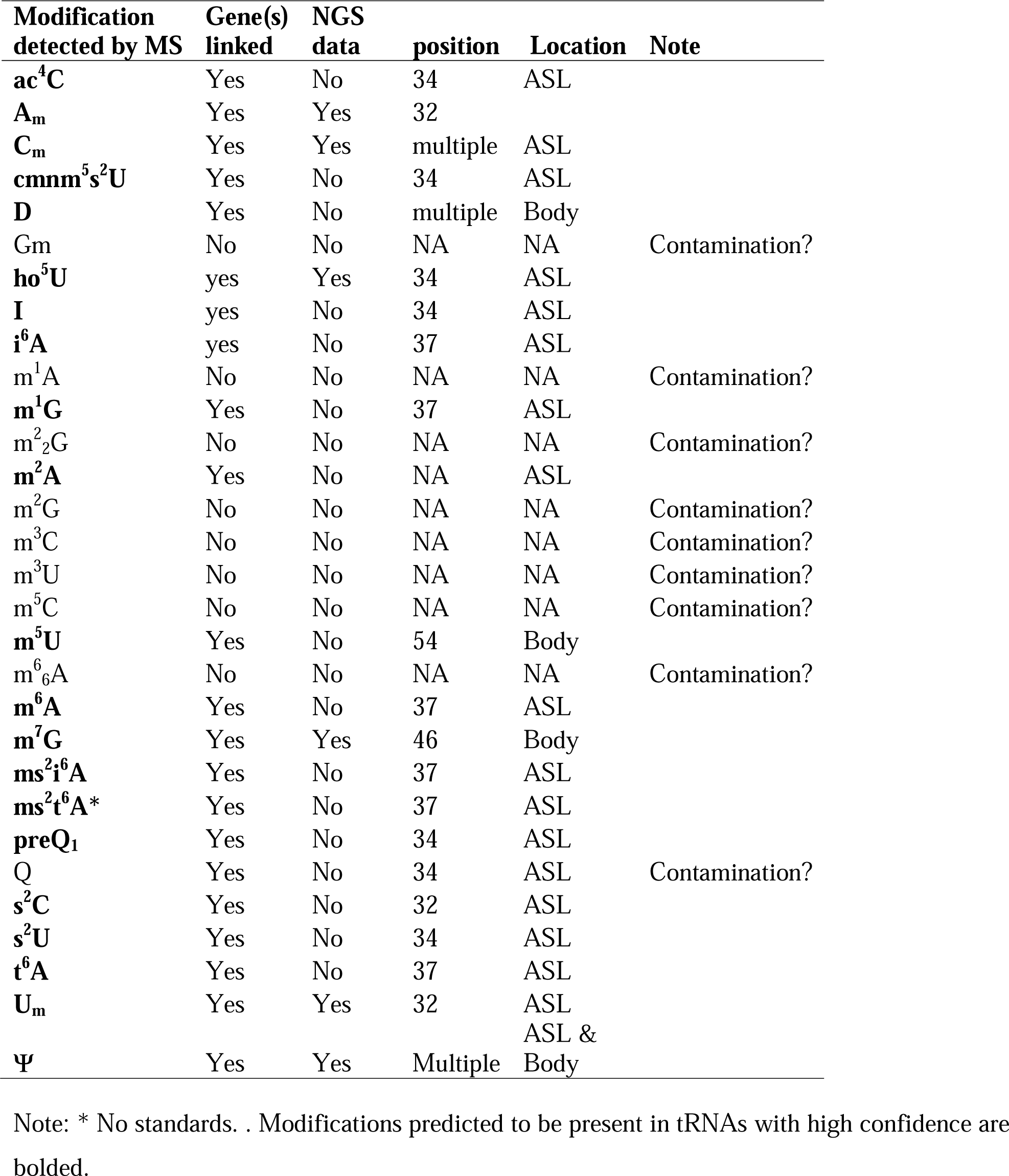
Summary of ribonucleosides identified by LC/MS in *B. henselae.*

### 3.2 Predicting the *B. henselae* tRNA Modification Gene Set

The compilation of tRNA modification genes, initially established for model organisms *E. coli* and *B. subtilis*^49^, underwent an update by incorporating two recently identified genes in *B. subtilis*^51^ into the dataset (**Table S5** and **S6**). This updated compilation served as a foundation for identifying tRNA modification genes encoded by *B. henselae* strain Houston 1, utilizing both BlastP and advanced search tools provided by BV-BRC and UniProt. The predictions generated were then cross-referenced with the modifications detected through LC-MS, and with the sequencing-based (RiboMethSeq, AlkAnilineSeq, and HydraPsiSeq) detection analyses results (**Fig. 2**), leading to the creation of the predicted tRNA modification map depicted in **Fig. 3**. In total, we predicted 26 genes participating in the insertion of 23 modifications (**Fig. 3** and **Table S7**).

**Figure 2.**
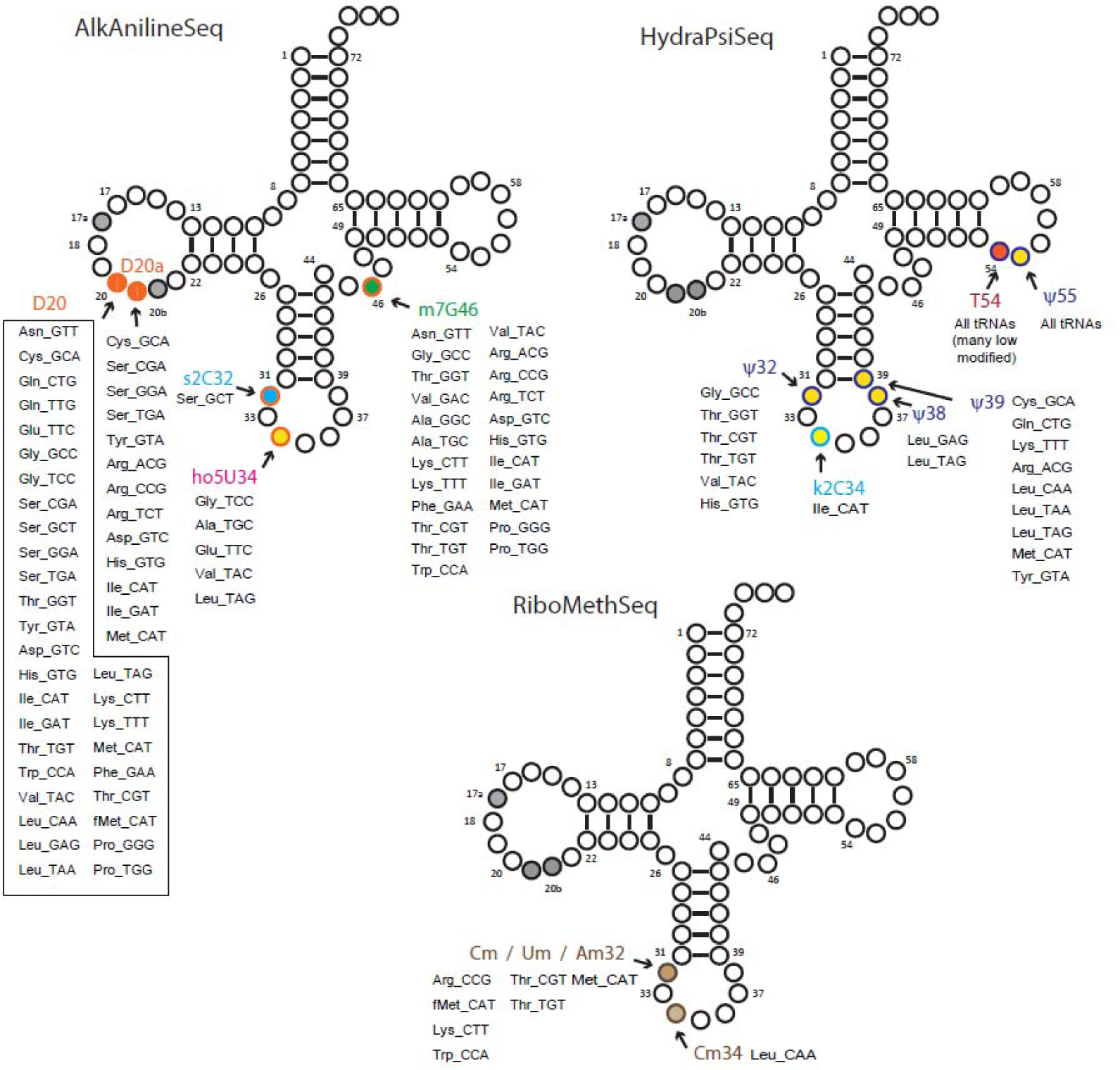
Overview of tRNA modifications mapping by deep sequencing-based protocols: AlkAnilineSeq, HydraPsiSeq and RiboMethSeq. Positions and identities of modified residue found in *B. henselae* tRNA species (shown by amino acid specificity and anticodon) are shown on a canonical cloverleaf tRNA structure. Long tRNA variable loops are not shown.

**Figure 3.**
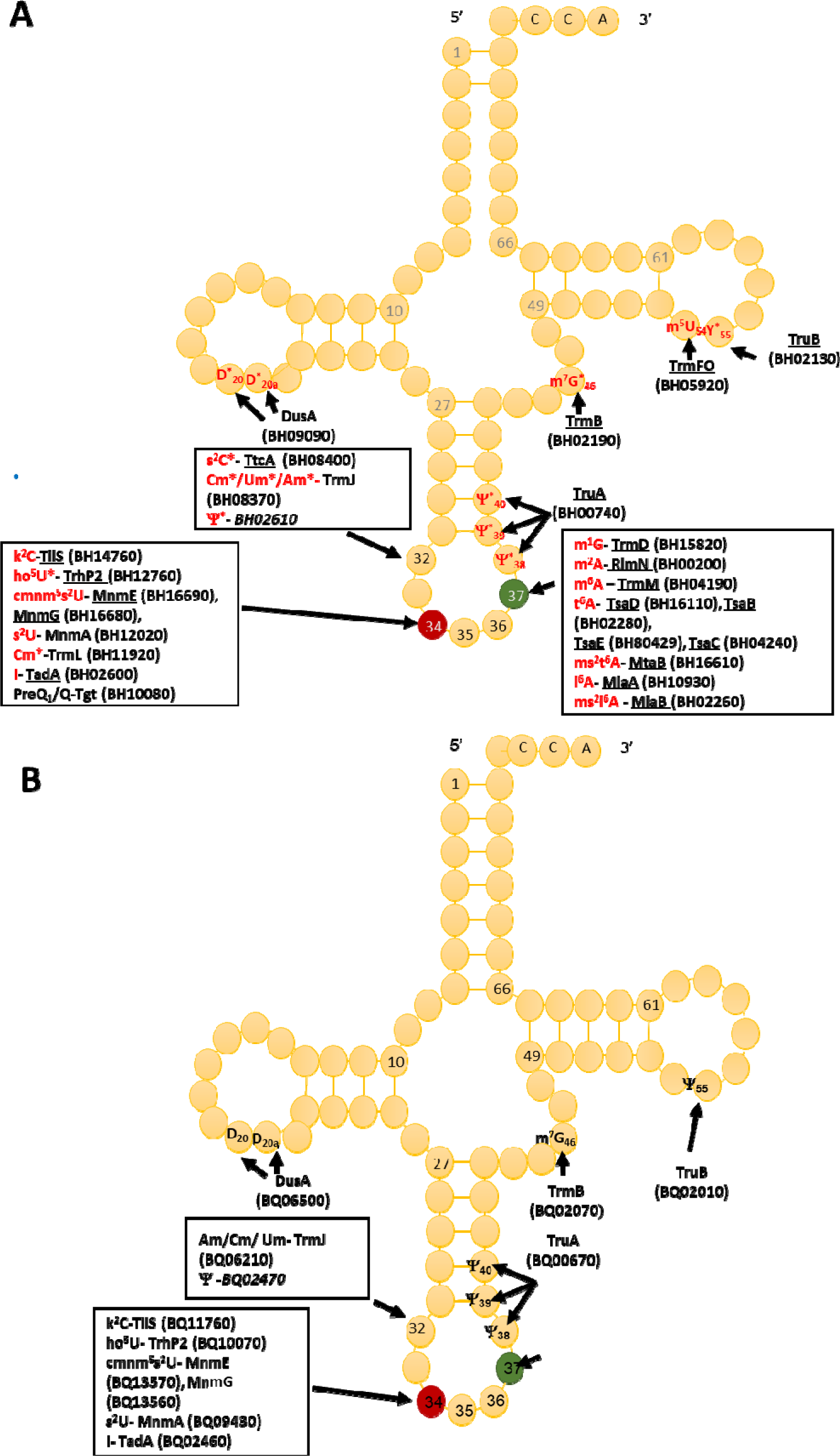
Prediction of *B. henselae* and *B. quintana* tRNA modifications maps and corresponding synthesis proteins. (A) *B. henselae*, underlined protein names denote high confidence based on orthology; in red are modifications detected by LC/MS in all samples. A “*” denotes that the modification at this position was confirmed by NGS. Locus tags are given in parentheses, and when italicized, denote a low-confidence prediction. (B) *B. quintana*, locus tags are given in parenthesis, and when italicized it denotes a low confidence.

For 18 of these modifications, the combination of gene information and LC/MS data yielded predictions with high confidence, as highlighted in (**Fig. 3 and Table 1).** The position of a few of those such as m^7^G46, ho^5^U34, and s^2^C32 were confirmed by sequencing **(Figs. 2 and S1)**. Additional mapping information was required for the remaining 5 modifications. Certain modification enzymes, like methyltransferases, pseudouridine synthases^52^ or dihydrouridine synthases^53^, have undergone multiple duplications and shifts in substrate specificity or may exhibit multisite specificity^54^. Consequently, predicting their specificities based solely on orthologous relationships can be challenging. Many bacteria possess multiple D modifications, introduced by different enzymes such as DusA/B/C in *E. coli* ^55^ or a single enzyme like DusB in *M. capricolum*^56^. In the *B. henselae* genome, the sole encoded member of the Dus family is a DusA homolog. While *E. coli*’s DusA modifies positions 20/20a^55^, we couldn’t exclude the possibility that in *B. henselae*, the enzyme had evolved a relatively broader specificity. However, AlkAnilineSeq analysis conclusively confirmed that the only positions modified by D in *Bartonella* tRNAs were 20/20a (and possibly 20b) (**Figs. 2 and S1**).

In the mapping of Ψ residues, the identification of homologs of TruA and TruB, and the absence of TruC and TruD, suggested modifications at positions 38-40 (TruA) and 55 (TruB) but excluding positions 13 and 65 formed by TruD and TruC, respectively. However, a conclusive determination for position 32, known to be modified in *E. coli* by a dual-specific RluA enzyme that modifies both rRNAs and tRNAs, was not as straightforward. Three predicted rRNA pseudouridine synthases are encoded in the *B. henselae* genome (BH02610, RluC/BH10200, or RluD/BH03820), and any of these might have shifted specificity to modify tRNAs, akin to the observed phenomenon with RluA and RluF in *E. coli*^57^. HydraPsiSeq analysis confirmed the absence of Ψ residues at positions 13 and 65 and the presence of Ψ at position 55 in many tRNAs and at positions 38/39/40 in a few tRNAs (**Figs. 2 and S2**) confirming the predictions based on the presence/absence of the corresponding synthesis genes. Ψ was also detected at position 32 in several tRNAs (**Figs. 2 and S2**). We propose that BH02610, the lone orphan pseudouridine synthase, catalyzes the formation of Ψ32 in *B. henselae*, but further experimental validation will be required. HydraPsiSeq protocol also detects 5’-modified U (namely m^5^U), which also shows protection against hydrazine cleavage, and k^2^C (lysidine) which is efficiently cleaved under conditions used in HydraPsiSeq (Fig. **S2**) The results of this mapping indicated that U54 in many tRNAs is only partially protected, indicating sub-stoichiometric modification by TrmFO. As anticipated, lysidine-related signal in HydraPsiSeq was found in tRNA^Ile^_CAU_ at wobble position C34 (**Fig. S2**). Finally, the presence of homologs of TrmJ and TrmL strongly predicts the presence of ribose −2’-O-methylations at positions 32 and 34, respectively, but the exact nature of the modified nucleosides could vary. RiboMethSeq analyses were able to map Am/Cm/Um at positions 32, but only Cm at position 34 (**Figs. 2 and S3**).

A significant number of modifications identified by LC/MS could not be attributed to specific genes (**Table 1**), including those present in substantial quantities like m^5^C (**Table S4a-b**). For m^5^C it was confirmed by bisulfite sequencing that indeed it is not present in the *B. henselae* tRNAs (data not shown). The levels of many of these “orphan” modifications exhibited considerable variation among the analyzed samples. For instance, m^2^_2_G was present in certain samples and absent in others **(Table S4a-b**). One plausible source for these modified nucleosides is rRNA from the bacteria or tRNA/rRNA from the host. Indeed, m^3^U, m^5^C, and m^6^_2_A are well-known rRNA modifications^33^, while contamination by mammalian host tRNAs, might contribute to the observed pools of m^1^A, G_m_, m^3^C, m^2^_2_G and m^2^G^33^.

### 3.3 Evolutionary Streamlining: Decoding Capacity Maintenance in *B. henselae* Through Modification Complexity Reduction

*B. henselae* exhibits a streamlined tRNA modification machinery, evident both in the reduced number of modifications and the simplicity of the existing ones. In comparison to *E. coli*, *B. henselae* encodes less than half the modification genes (26 genes compared to *E. coli*’s 59). Studies on insect symbionts have indicated that modifications of the tRNA body are the first to diminish and can even be entirely lost by reductive evolution, as observed in the genome of louse symbionts^58^. Notably, common tRNA body modifications such as D16/17, Ψ13/65, and s^4^U8 are absent in *B. henselae* tRNAs based on the absence of the genes and confirmed by the lack of the NGS signals (data not shown). DusB family is predicted to be the ancestral Dus enzyme ^59^, and depending on the organism it modifies position 16 or is multisite-specific^60^. It seems to have been lost near the root of the *Bartonellaceae* clade (**Fig. S4B**). The only remaining Dus enzyme in *B. henselae* is DusA which is found in all *Hyphomicrobiales* (**Fig. S4A**). TruC and TruD and ThiI are totally absent in *Alphaprotebacteria*, hence the observed losses in the *Bartonellaceae* are not recent events **(Fig. S5).**

Losing modifications in the Anticodon Stem Loop (ASL) should significantly impact decoding capacity, efficiency, and accuracy. Only two ASL modifications found in *E. coli* are absent in *B. henselae*: ac^4^C34 and Ψ35. The first is found in the elongator tRNA^Met^_CAU_ and prevents misreading of the near cognate AUA (Ile) codon^61^. The second affects the decoding efficiency of Tyr codon stretches^57^. However, as the synthesis enzymes for both these modifications (TmcA and RluF) seem to be absent in nearly all *Alphaproteobacteria* (**Fig. S5**), organisms must compensate for the absence of these modifications.

Interestingly, several complex modifications present in *E. coli* exist in *B. henselae* but in a simpler form that requires a truncated synthesis pathway with fewer genes. The simplification of the Q pathway to salvaging and inserting preQ_1_ and possibly q bases was reported previously^26^, but it looks like all complex pathways have been simplified in this organism. For instance, the cmnm^5^s^2^U precursor, synthesized by MmmGEA enzymes, is present, but its derivative mnm^5^s^2^U, requiring two additional enzymatic steps, is not. Similarly, the ho^5^U precursor is present, but not its derivative, cmo^5^U. Likewise, the t^6^A modification is present but it is not cyclized to ct^6^A nor further modified to m^6^t^6^A. These simplifications of complex ASL modifications to minimal forms have facilitated the reduction of the number of tRNA modification genes without affecting the capacity of the 43 *B. henselae* tRNAs to decode the full set of codons (**Fig. 1**). There is only one example of a modification that is more complex in *B. henselae* than in *E. coli*: ms^2^t^6^A. The enzyme involved in its synthesis, MtaB, is widespread in *Alphaprotebacteria*, so its presence in *Bartonellaceae* reflects the evolutionary history of the species (**Fig. S5**).

### 3.4 Predicting the tRNA modification gene set in *B. quintana* shows further reductions

A parallel analysis was executed to predict the tRNA modification gene sets in *B. quintana*, as depicted in **Fig. 2B** and detailed in **Table S8**. This analysis identified 22 genes linked to 23 modifications. While most genes were found in both *Bartonella* species, instances of gene decay were observed, resulting in the loss of corresponding modifications in the extracted tRNA digests (**Table S4a-b and Fig. 4)**. Prior findings had noted the decay of the *tgt* gene^26^, and three more instances were identified: *ttcA*, *trmFO*, and *trmL*. In each case, the complete gene is present in *B. henselae*, but only fragments are discernible at the corresponding loci in the sequenced *B. quintana* Toulouse genome (**Fig. 5**). Notably, the decay of *tgt*, *ttcA*, and *trmFO* genes appears consistent across all sequenced *B. quintana* strains, while the truncation of *trm*L seems specific to the *B. quintana* Toulouse strain. Results from LC/MS/MS analyses of bulk tRNAs extracted from *B. quintana* seem to align with these losses. For instance, the disappearance of the *ttcA* gene correlates with the absence of s^2^C in the nucleoside analysis profile from *B. quintana*, contrasting with its presence in *B. henselae* (**Fig. 4** and **Table S4a-b**). Levels of C_m_ are not dramatically affected in *B. quintana* (**Fig. 4** and **Table S4a-b**). Given that C_m_ can also be derived from contaminating rRNAs or host tRNAs and that RiboMethSeq for *B. quintana* tRNAs was not performed, additional targeted experiments will be essential to validate the absence of this modification at position 34. Members of the TtcA family are present in most *Bartonella* species (**Fig. S4B**). s^2^C32 has been shown to have a role in preventing frameshift^62^ but is also important to restrict the ability of tRNA^Arg^ to decode the rare arginine codon CGA^63^. Losing the Cm methylation in tRNA^Leu^_CAA_, reduces the efficiency of codon-wobble base interaction, as demonstrated in an amber suppressor system^64^. It is therefore expected that a loss of translation accuracy should occur with the loss of both C_m_ and s^2^C modifications in *B. quintana*, a phenomenon often observed in host-restricted organisms^65^.

**Figure 4.**
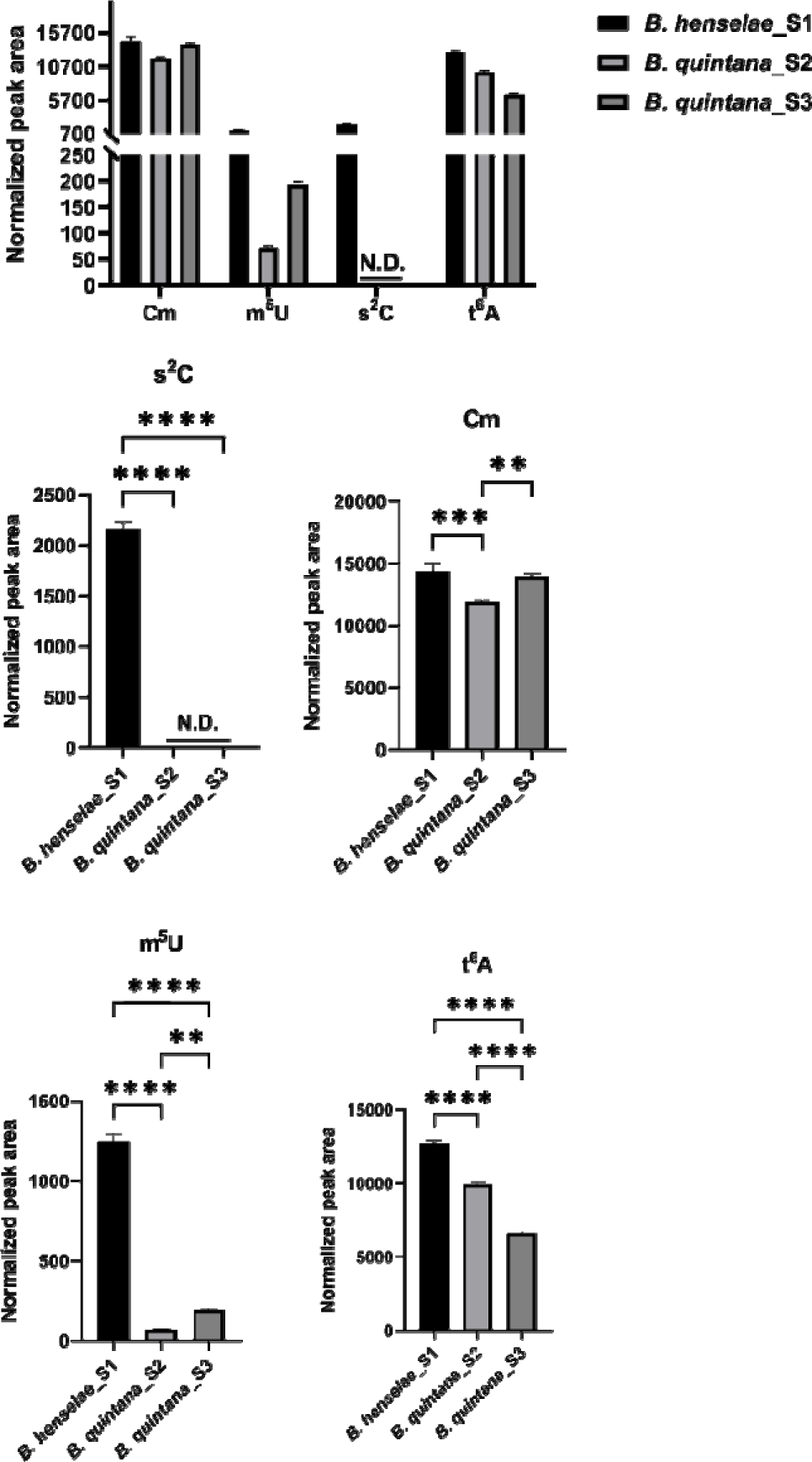
Normalized peak area values of Cm, m^5^U, s^2^C and t^6^A modifications in *B. henselae* and *B. quintana*. 600 ng of hydrolysate was injected. For *B. quintana*, 2 biological replicates were provided. Three technical replicates were performed for each sample. The signals were confirmed by both qualifier and quantifier transitions. Data represent the means ±SD for all replicates. Statistical analysis was done by one-way analysis of variance (ANOVA) with Prism 9, and only comparisons with P values less than or equal to 0.05; P <0.05, P <0.01, P <0.001 and P <0.0001 are denoted as *, **, *** and **** respectively. All modifications were validated with standards. N.D.: Not detected.

**Figure 5.**
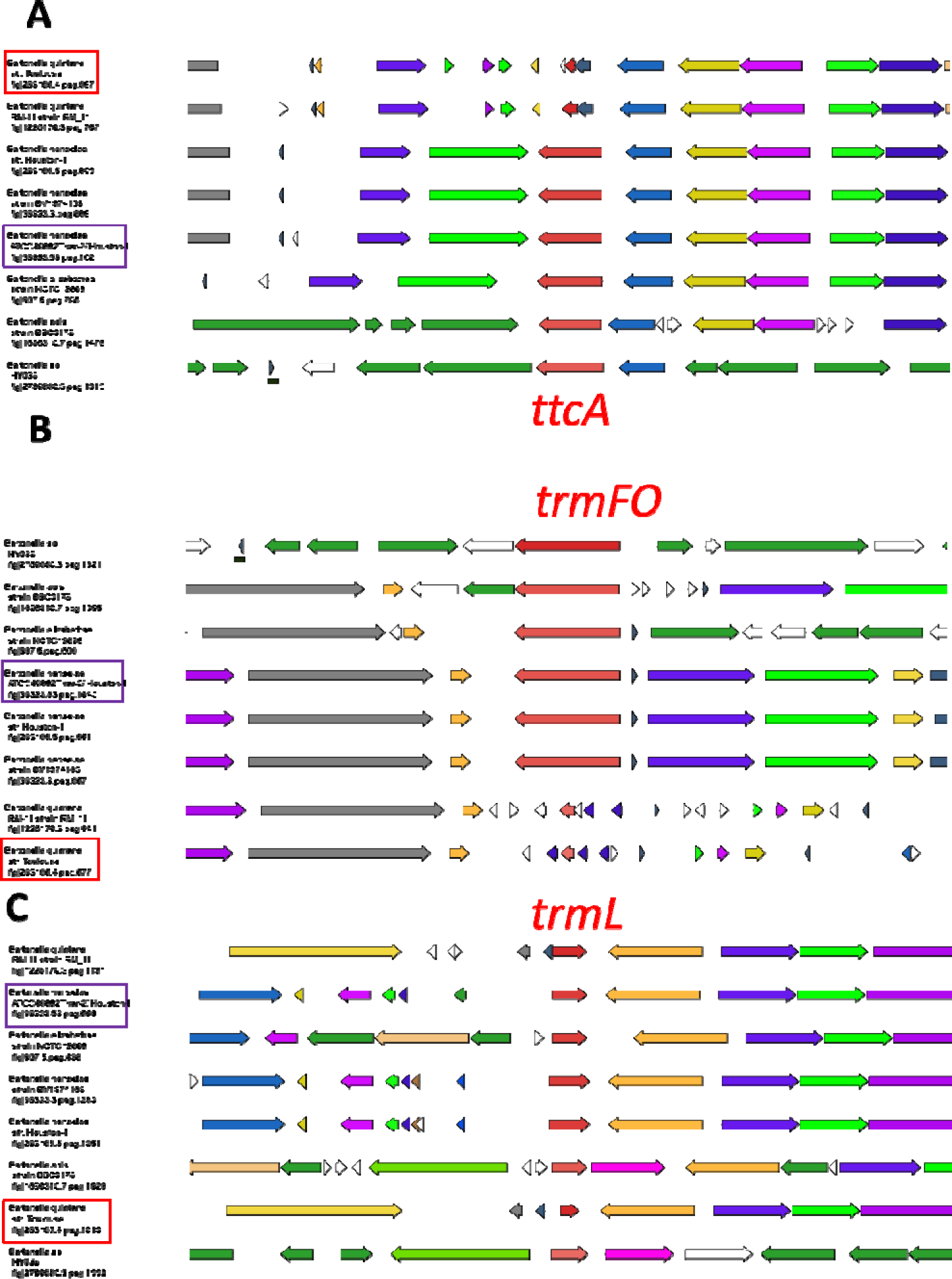
Decay of *ttcA*, *trmL* and *trmFO genes in B. quintana* species. Gene neighborhoods surrounding the three modificationgenes in *Bartonella* species. *B. quintana* str. Toulouse and the *B. henselae* Houston-I are boxed in red and purple, respectively. The BV-BRC gene ids (or fig ids) of the specific anchoring genes [*ttcA* in (A), *trmFO* in (B) and *trmL* in (C)] are given for every cluster.

Finally, m^5^U levels are reduced by approximately 80% in *B. quintana* compared to *B. henselae* (**Fig. 4** and **Table S4a-b**). The residual m^5^U could be derived from contaminating rRNA and/or host tRNAs^33^. Two non-orthologous families of methylases can catalyze the formation of m^5^U54 in tRNA: the SAM-dependent TrmA and the FAD-dependent TrmFO. The two enzymes are mutually exclusive, and genomes encode one or the other^66^. Members of the alphaproteobacterial clade use TrmFO (**Fig. S5**), hence its presence in *Bartonellaceae* is to be expected. The loss of m^5^U54 in *B. quintana* is like what is observed in other organisms with minimal genomes ^46,58^.

## 4. Conclusions

This study shows that the tRNA modifications profiles and most corresponding genes can be predicted for the facultative intracellular pathogens *B. henselae* and *B. quintana*, but one must be wary of contamination by host tRNAs and rRNA and combine different types of evidence. Also, it is possible that certain modifications may have been missed in the study, e.g., /if novel pathways are present to insert modifications that could not be chemically identified because of a lack of standards. Nevertheless, even with these potential omissions, it is possible to conclude that while *B. henselae* has not greatly reduced the number of tRNA isoacceptors, they are all in single copy. In addition, the number of modified bases in tRNAs has been reduced from 43 in *E. coli* to 28 in *B. henselae* (a ∼35% relative reduction). Finally, the number of tRNA modification genes has undergone an even greater reduction with 59 genes in *E. coli* to 26 in *B. henselae* (a ∼56% relative reduction) because of the simplification of the most complex pathways. Further simplifications of the tRNA modifications apparatus are observed in *B. quintana* with an additional loss of four modifications (preQ_1_, C_m_. s^2^C and m^5^U) compared to *B*. *henselae*, which correlates with the extensive genome reduction following its divergence from *B. henselae* and specialization for the human host and louse vector ^22^.

## Supporting information

Supplemental_Tables_S1toS11

Supplemental_Figures_S1toS5

## Acknowledgments

This work was funded by the National Institutes of Health (project numbers GM070641, ES026856, and ES024615) and the National Research Foundation of Singapore through the Singapore-MIT Alliance for Research and Technology Antimicrobial Resistance Interdisciplinary Research Group.

## References

1. Zaher HS, Green RF. Fidelity at the molecular level: lessons from protein synthesis. Cell. 2009 Feb 20;136(4):746–762.

2. Berg MD, Brandl CJ. Transfer RNAs: Diversity in form and function. RNA Biology. 2021;18(3):316–339.

3. El Yacoubi B, Bailly M, de Crécy-Lagard V. Biosynthesis and function of posttranscriptional modifications of transfer RNAs. Annu Rev Genet. 2012;46(1):69–95.

4. Chan CTY, Dyavaiah M, DeMott MS, Taghizadeh K, Dedon PC, Begley TJ. A quantitative systems approach reveals dynamic control of tRNA modifications during cellular stress. PLoS Genet. 2010;6(12):1–9.

5. de Crécy-Lagard V, Jaroch M. Functions of bacterial tRNA modifications: from ubiquity to diversity. Trends Microbiol. 2021 Jan 1;29(1):41–53.

6. Wang L, Lin S. Emerging functions of tRNA modifications in mRNA translation and diseases. J Genet Genomics. 2023 Apr 1;50(4):223–232.

7. Helm M, Alfonzo JD. Posttranscriptional RNA modifications: playing metabolic games in a cell’s chemical legoland. Chem Biol. 2014;21(2):174–185.

8. Persson BC. Modification of tRNA as a regulatory device. Mol Microbiol. 1993;8(6):1011–1016.

9. Koh CS, Sarin LP. Transfer RNA modification and infection – implications for pathogenicity and host responses. Biochim Biophys Acta - Gene Regul Mech. 2018;1861(4):419–432.

10. Fruchard L, Babosan A, Carvalho A, Lang M, Li B, Duchateau M, Giai-Gianetto Q, Matondo M, Bonhomme F, Fabret C, Namy O, de Crécy-Lagard V, Mazel D, Baharoglu Z. Queuosine modification of tRNA-tyrosine elicits translational reprogramming and enhances growth of *Vibrio cholerae* with aminoglycosides. bioRxiv. 2022 Sep 26;2022.09.26.509455.

11. Antoine L, Bahena-Ceron R, Devi Bunwaree H, Gobry M, Loegler V, Romby P, Marzi S. RNA modifications in pathogenic bacteria: impact on host adaptation and virulence. genes. 2021; 12(8):1125.

12. Kimura S, Dedon PC, Waldor MK. Comparative tRNA sequencing and RNA mass spectrometry for surveying tRNA modifications. Nat Chem Biol. 2020;16(9):964–972.

13. Tomasi, F. G.; Kimura, S.; Rubin, E. J.; Waldor, M. K. A tRNA Modification in *Mycobacterium tuberculosis* facilitates optimal intracellular growth. Elife. 2023 Sep 27;12:RP87146

14. Koshla O, Vogt LM, Rydkin O, Sehin Y, Ostash I, Helm M, Ostash B. Landscape of post-transcriptional tRNA modifications in *Streptomyces albidoflavus* J1074 as portrayed by mass spectrometry and genomic data mining. J Bacteriol. 2023;205(1).

15. Loterio RK, Zamboni DS, Newton HJ. Keeping the host alive – lessons from obligate intracellular bacterial pathogens. Pathog Dis. 2021 Dec;79(9): ftab052.

16. McClure EE, Chávez ASO, Shaw DK, Carlyon JA, Ganta RR, Noh SM, Wood DO, Bavoil PM, Brayton KA, Martinez JJ, McBride JW, Valdivia RH, Munderloh UG, Pedra JHF. Engineering of obligate intracellular bacteria: progress, challenges and paradigms. Nat Rev Microbiol. 2017;15:544–558.

17. Albalat, R.; Cañestro, C. Evolution by Gene Loss. Nature Reviews Genetics. 2016 Jul;17(7):379–91

18. Andersson SGE, Kurland CG. Reductive evolution of resident genomes. Trends Microbiol. 1998;6(7):263–268.

19. Rovid Spickler, A. Cat Scratch Disease; 2005. www.cfsph.iastate.edu.

20. Raoult D. From cat scratch disease to *Bartonella henselae* infection. Clin Infect Dis. 2007 Dec 15;45(12):1541–1542.

21. Foucault C, Brouqui P, Raoult D. *Bartonella quintana* characteristics and clinical management. Emerg Infect Dis. 2006 Feb;12(2):217–223.

22. Alsmark CM, Frank AC, Karlberg EO, Legault BA, Ardell DH, Canbäck B, Eriksson AS, Näslund AK, Handley SA, Huvet M, La Scola B, Holmberg M, Andersson SGE. The louse-borne human pathogen *Bartonella quintana* is a genomic derivative of the zoonotic agent *Bartonella henselae*. Proc Natl Acad Sci U S A. 2004 Jun 29;101(26):9716–9721.

23. Segers FH, Kešnerová L, Kosoy M, Engel P. Genomic changes associated with the evolutionary transition of an insect gut symbiont into a blood-borne pathogen. ISME J. 2017 May;11(5):1232–1244.

24. Engel P, Dehio C. Genomics of host-restricted pathogens of the genus *Bartonella*. Genome Dyn. 2009;6: 158–169.

25. Battisti JM, Minnick MF. Laboratory maintenance of *Bartonella quintana*. Curr Protoc Microbiol. 2008 May;10(1):3C.1.1-3C.1.13.

26. Quaiyum S, Yuan Y, Sun G, Ratnayakec RMM, Hutinet G, Dedon PC, Minnick MF, de Crécy-Lagard V. Queuosine salvage in *Bartonella henselae* Houston 1: A unique evolutionary path. bioRxiv. 2023 Dec 5;2023.12.05.570228.

27. Chan PP, Lowe TM. GtRNAdb 2.0: An expanded database of transfer RNA genes identified in complete and draft genomes. Nucleic Acids Res. 2016 Jan 4;44(D1): D184–D189.

28. Olson RD, Assaf R, Brettin T, Conrad N, Cucinell C, Davis JJ, Dempsey DM, Dickerman A, Dietrich EM, Kenyon RW, Kuscuoglu M, Lefkowitz EJ, Lu J, Machi D, Macken C, Mao C, Niewiadomska A, Nguyen M, Olsen GJ, Overbeek JC, Parrello B, Parrello V, Porter JS, Pusch GD, Shukla M, Singh I, Stewart L, Tan G, Thomas C, VanOeffelen M, Vonstein V, Wallace ZS, Warren AS, Wattam AR, Xia F, Yoo H, Zhang Y, Zmasek CM, Scheuermann RH, Stevens RL. Introducing the bacterial and viral bioinformatics resource center (BV-BRC): A resource combining PATRIC, IRD, and ViPR. Nucleic Acids Res. 2023 Jan 4;51(1D):D678–D689.

29. Kanehisa M, Furumichi M, Sato Y, Ishiguro-Watanabe M, Tanabe M. KEGG: Integrating viruses and cellular organisms. Nucleic Acids Res. 2021 Jan 8;49(D1):D545–D551.

30. Bateman A, Martin M-J, Orchard S, Magrane M, Ahmad S, Alpi E, Bowler-Barnett EH, Britto R, Bye-A-Jee H, Cukura A, Denny P, Dogan T, Ebenezer T, Fan J, Garmiri P, da Costa Gonzales LJ, Hatton-Ellis E, Hussein A, Ignatchenko A, Insana G, Ishtiaq R, Joshi V, Jyothi D, Kandasaamy S, Lock A, Luciani A, Lugaric M, Luo J, Lussi Y, MacDougall A, Madeira F, Mahmoudy M, Mishra A, Moulang K, Nightingale A, Pundir S, Qi G, Raj S, Raposo P, Rice DL, Saidi R, Santos R, Speretta E, Stephenson J, Totoo P, Turner E, Tyagi N, Vasudev P, Warner K, Watkins X, Zaru R, Zellner H, Bridge AJ, Aimo L, Argoud-Puy G, Auchincloss AH, Axelsen KB, Bansal P, Baratin D, Batista Neto TM, Blatter M-C, Bolleman JT, Boutet E, Breuza L, Gil BC, Casals-Casas C, Echioukh KC, Coudert E, Cuche B, de Castro E, Estreicher A, Famiglietti ML, Feuermann M, Gasteiger E, Gaudet P, Gehant S, Gerritsen V, Gos A, Gruaz N, Hulo C, Hyka-Nouspikel N, Jungo F, Kerhornou A, Le Mercier P, Lieberherr D, Masson P, Morgat A, Muthukrishnan V, Paesano S, Pedruzzi I, Pilbout S, Pourcel L, Poux S, Pozzato M, Pruess M, Redaschi N, Rivoire C, Sigrist1 CJA, Sonesson K, Sundaram S, Wu CH, Arighi CN, Arminski L, Chen C, Chen Y, Huang H, Laiho K, McGarvey P, Natale DA, Ross K, Vinayaka CR, Wang Q, Wang Y, Zhang J. UniProt: The universal protein knowledgebase in 2023. Nucleic Acids Res. 2023 Jan 4;51(D1):D523-D531.

31. Altschul SF, Madden TL, Schaffer AA, Zhang J, Zhang Z, Miller W, Lipman DJ. Gapped BLAST and PSI-BLAST: A new generation of protein database search programs. Nucleic Acids Res. 1997 Sep 1;25(17):3389–3402.

32. NCBI Resource Coordinators. Database resources of the national center for biotechnology information. Nucleic Acids Res. 2016 Jan 4;44: D7–D19.

33. Boccaletto P, MacHnicka MA, Purta E, Pitkowski P, Baginski B, Wirecki TK, de Crécy-Lagard V, Ross R, Limbach PA, Kotter A, Helm M, Bujnicki JM. MODOMICS: A database of RNA modification pathways. 2017 update. Nucleic Acids Res. 2018 Jan 4;46(D1): D303-D307.

34. Letunic I, Bork P. Interactive Tree of Life (ITOL) v5: An online tool for phylogenetic tree display and annotation. Nucleic Acids Res. 2021 Jul 2;49(W1): W293–W29.

35. Marchand V, Pichot F, Thüring K, Ayadi L, Freund I, Dalpke A, Helm M, Motorin Y. Next-generation sequencing-based RibOxi-MeRIP-seq protocol for analysis of tRNA 2’-O-methylation. Biomolecules. 2017 Jan 16;7(1):13.

36. Marchand V, Blanloeil-Oillo F, Helm M, Motorin Y. Illumina-based RiboMethSeq approach for mapping of 2’-O-Me residues in RNA. Nucleic Acids Res. 2016 Aug 19;44(16): e135.

37. Werner S, Schmidt L, Marchand V, Kemmer T, Falschlunger C, Sednev MV, Bec G, Ennifar E, Höbartner C, Micura R, Motorin Y, Hildebrandt A, Helm M. Machine learning of reverse transcription signatures of variegated polymerases allows mapping and discrimination of methylated purines in limited transcriptomes. Nucleic Acids Res. 2020 Apr 6;48(7):3734–3746.

38. Motorin Y, Marchand V. Detection and analysis of RNA ribose 2’-O-methylations: challenges and solutions. Genes (Basel). 2018 Dec 4;9(12):642.

39. Thorvaldsdóttir H, Robinson JT, Mesirov JP. Integrative genomics viewer (IGV): high-performance genomics data visualization and exploration. Brief Bioinform. 2013 Mar;14(2):178–192.

40. Marchand V, Ayadi L, Ernst FGM, Hertler J, Bourguignon-Igel V, Galvanin A, Kotter A, Helm M, Lafontaine DLJ, Motorin Y. AlkAniline-Seq: Profiling of m^7^G and m^3^C RNA modifications at single nucleotide resolution. Angew Chem Int Ed Engl. 2018 Dec 14;57(51):16785–16790.

41. Marchand V, Bourguignon-Igel V, Helm M, Motorin Y. Mapping of 7-Methylguanosine (m^7^G), 3-Methylcytidine (m^3^C), Dihydrouridine (D), and 5-Hydroxycytidine (Ho5C) RNA modifications by AlkAniline-Seq. In: Methods in Enzymology; 2021; 658:25-47.

42. de Brouwer APM, Abou Jamra R, Körtel N, Soyris C, Polla DL, Safra M, Zisso A, Powell CA, Rebelo-Guiomar P, Dinges N, Morin V, Stock M, Hussain M, Shahzad M, Riazuddin S, Ahmed ZM, Pfundt R, Schwarz F, de Boer L, Reis A, Grozeva D, Raymond FL, Riazuddin S, Koolen DA, Minczuk M, Roignant J-Y, van Bokhoven H, Schwartz S. Variants in PUS7 cause intellectual disability with speech delay, microcephaly, short stature, and aggressive behavior. Am J Hum Genet. 2018 Dec 6;103(6):1045–1052.

43. Behm-Ansmant I, Helm M, Motorin Y. Use of specific chemical reagents for detection of modified nucleotides in RNA. J Nucleic Acids. 2011; 2011:408053.

44. Motorin Y, Muller S, Behm-Ansmant I, Branlant C. Identification of modified residues in RNAs by reverse transcription-based methods. Methods Enzymol. 2007; 425:21–53.

45. Schaefer M, Pollex T, Hanna K, Lyko F. RNA cytosine methylation analysis by bisulfite sequencing. Nucleic Acids Res. 2009;37(2): e12.

46. Grosjean H, Breton M, Sirand-Pugnet P, Tardy F, Thiaucourt F, Citti C, Barré A, Yoshizawa S, Fourmy D, de Crécy-Lagard V, Blanchard A. Predicting the minimal translation apparatus: lessons from the reductive evolution of Mollicutes. PLoS Genet. 2014; 10(5): e1004363.

47. Chan PP, Lowe TM. GtRNAdb 2.0: An expanded database of transfer RNA genes identified in complete and draft genomes. Nucleic Acids Res. 2016 Jan;44(D1): D184–D189.

48. Grosjean H, de Crécy-Lagard V, Marck C. Deciphering synonymous codons in the three domains of life: co-evolution with specific tRNA modification enzymes. FEBS Lett. 2010 Jan 21;584(2):252.

49. de Crécy-Lagard V, Ross RL, Jaroch M, Marchand V, Eisenhart C, Brégeon D, Motorin Y, Limbach PA. Survey and validation of tRNA modifications and their corresponding genes in *Bacillus subtilis* strain 168. Biomolecules. 2020 Jul;10(7):1–23.

50. Grosjean H, Westhof E. An integrated, structure-and energy-based view of the genetic code. Nucleic Acids Res. 2016 Sep 30;44(17):8020–8040.

51. Jaroch M, Sun G, Tsui HC, Reed C, Sun J, Jörg M, Winkler ME, Rice KC, Stich TA, Dedon PC, Santos PC Dos, de Crécy-Lagard V. Alternate routes to mnm5s2U synthesis in Gram-positive bacteria. bioRxiv. 2023 Dec 21;2023.12.21.572861.

52. Hamma T, Ferre-D’Amare AR. Pseudouridine synthases. Chem Biol. 2006 Nov;13(11):1125–1135.

53. Bou-Nader C, Montémont H, Guérineau V, Jean-Jean O, Brégeon D, Hamdane D. Unveiling structural and functional divergences of bacterial tRNA dihydrouridine synthases: perspectives on the evolution scenario. Nucleic Acids Res. 2018 Feb 16;46(3):1386–1394.

54. Barraud P, Tisné C. To be or not to be modified: Miscellaneous aspects influencing nucleotide modifications in tRNAs. IUBMB Life. 2019 Aug;71(8):1126–1140.

55. Bishop AC, Xu J, Johnson RC, Schimmel P, de Crécy-Lagard V. Identification of the tRNA-dihydrouridine synthase family. J Biol Chem. 2002 Jul 12;277(28):25090–25095.

56. Faivre B, Lombard M, Fakroun S, Vo CDT, Goyenvalle C, Guérineau V, Pecqueur L, Fontecave M, de Crécy-Lagard V, Brégeon D, Hamdane D. Dihydrouridine synthesis in tRNAs is under reductive evolution in Mollicutes. RNA Biol. 2021;18 (12):1–12.

57. Addepalli B, Limbach PA. Pseudouridine in the anticodon of *Escherichia coli* tRNA^Tyr^(QΨA) is catalyzed by the dual specificity enzyme RluF. J Biol Chem. 2016 Oct 14;291(42):22327–22337.

58. de Crécy-Lagard V, Marck C, Grosjean H. Decoding in *Candidatus Riesia pediculicola*, close to a minimal tRNA modification set? Trends Cell Mol Biol. 2012; 7:11–34.

59. Bou-Nader C, Montémont H, Guérineau V, Jean-Jean O, Brégeon D, Hamdane D. Unveiling structural and functional divergences of bacterial tRNA dihydrouridine synthases: perspectives on the evolution scenario. Nucleic Acids Res. 2018;46(3):1386–1394.

60. Faivre B, Lombard M, Fakroun S, Vo CDT, Goyenvalle C, Guérineau V, Pecqueur L, Fontecave M, de Crécy-Lagard V, Brégeon D, Hamdane D. Dihydrouridine synthesis in tRNAs is under reductive evolution in Mollicutes. RNA Biol. 2021;18(12).

61. Kawai G, Hashizume T, Miyazawa T, McCloskey JA, Yokoyama S. Conformational characteristics of 4-acetylcytidine found in tRNA. Nucleic Acids Symp Ser. 1989;21:61–62.

62. Jager G, Leipuviene R, Pollard MG, Qian Q, Björk GR. The conserved Cys-X1-X2-Cys motif present in the TtcA protein is required for the thiolation of cytidine in position 32 of tRNA from *Salmonella enterica* serovar *Typhimurium*. J Bacteriol. 2004;186(3):750–7.

63. Vangaveti S, Cantara WA, Spears JL, DeMirci H, Murphy FV, Ranganathan SV, Sarachan KL, Agris PF. A structural basis for restricted codon recognition mediated by 2-thiocytidine in tRNA containing a wobble position inosine. J Mol Biol. 2020;432(4): 913–929.

64. Benítez-Páez A, Villarroya M, Douthwaite S, Gabaldón T, Armengod ME. YibK is the 2’-O-methyltransferase TrmL that modifies the wobble nucleotide in *Escherichia coli* tRN^Aleu^ isoacceptors. RNA. 2010;16(11):2131–2143.

65. Melnikov SV, van den Elzen A, Stevens DL, Thoreen CC, Söll D. Loss of protein synthesis quality control in host-restricted organisms. Proc Natl Acad Sci U S A. 2018;115(49): E11505–E11512.

66. Myllykallio H, Sournia P, Heliou A, Liebl U. Unique features and anti-microbial targeting of folate-and flavin-dependent methyltransferases required for accurate maintenance of genetic information. Front Microbiol. 2018; 9:918.

