## Supplemental_Figures_S1toS5 for "Mapping the tRNA Modification Landscape of *Bartonella henselae* Houston I and *Bartonella quintana* Toulouse"

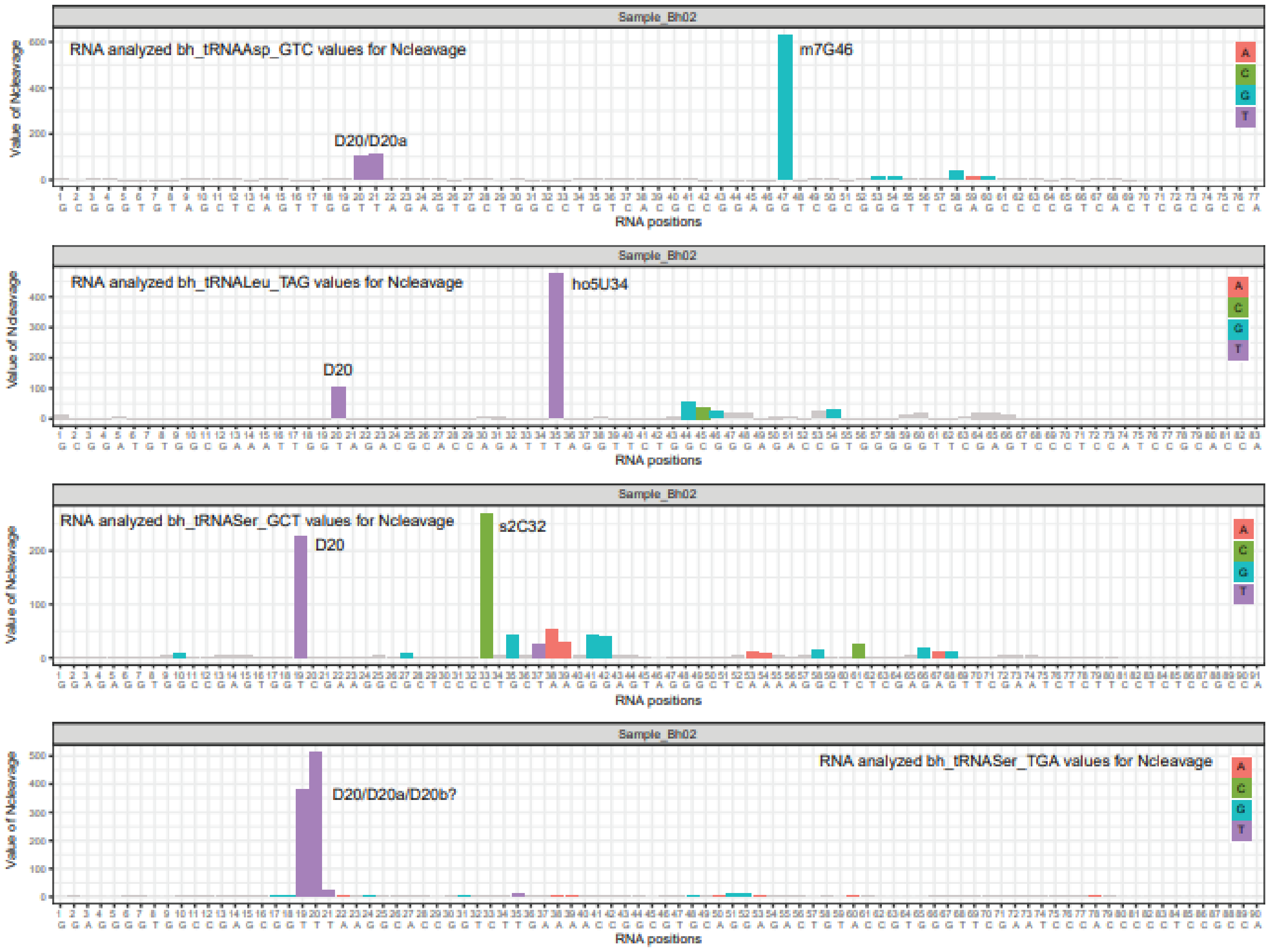


**Figure S1.** Representative AlkAnilineSeq signals observed in *B. henselae* tRNAs. Nucleotides are color-coded according to their identity. The sequence of tRNA with consecutive numbering of tRNA nucleotides is shown at the bottom. Traces are shown only for the Normalized cleavage (Ncleavage) score, which corresponds to the proportion of reads mapped to a given position divided by the total number of reads mapped to a given RNA. This score provides high specificity but is less sensitive than other AlkAnilineSeq scores. Signals for D20/D20a and m7G46 in tRNAAsp_GTC (top trace), signals for D20 and presumable ho5U34 in tRNALeu_TAG (middle top), D20 and s2C34 in tRNASer_GCT (middle bottom) and small signal for presumable D20b in tRNASer_TGA (bottom trace) are shown.

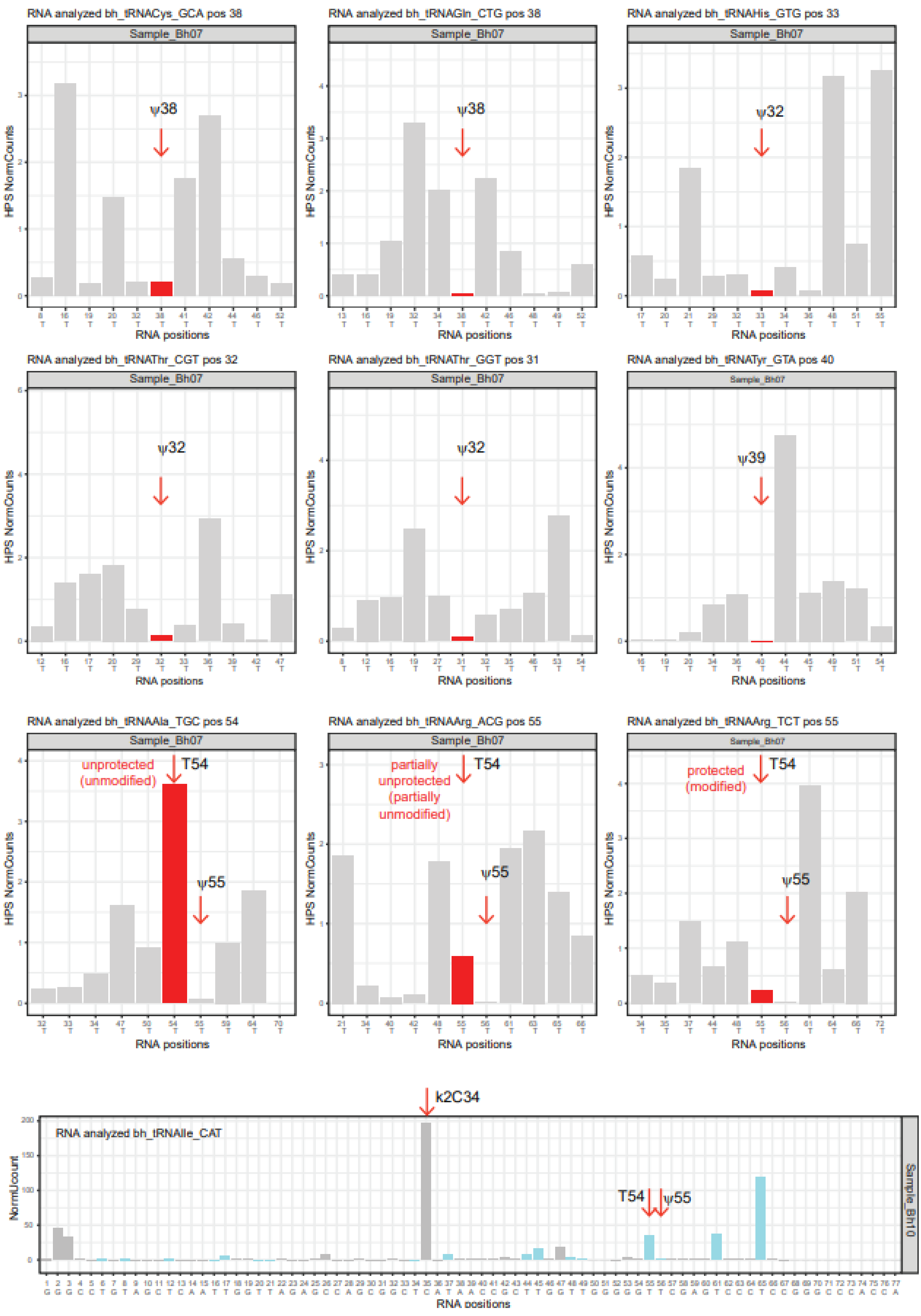


**Figure S2.** Mapping of protected pseudouridine (ψ) and $m^5U$(rT)54 residues in *B. henselae* tRNAs. Normalized hydrazine cleavage at 11 consecutive U residues (NormUcount) is shown for different protected (modified) positions, centered and red-colored. Partially protected $m^5U$(rT)54 residues are shown in the bottom traces, while ψ55 is highly protected in all tRNAs. The full profile for tRNAIle_CAT showing strong cleavage at $k^2C34$ is shown at the bottom. The sequence of tRNAIle_CAT with consecutive numbering of tRNA nucleotides is shown at the bottom

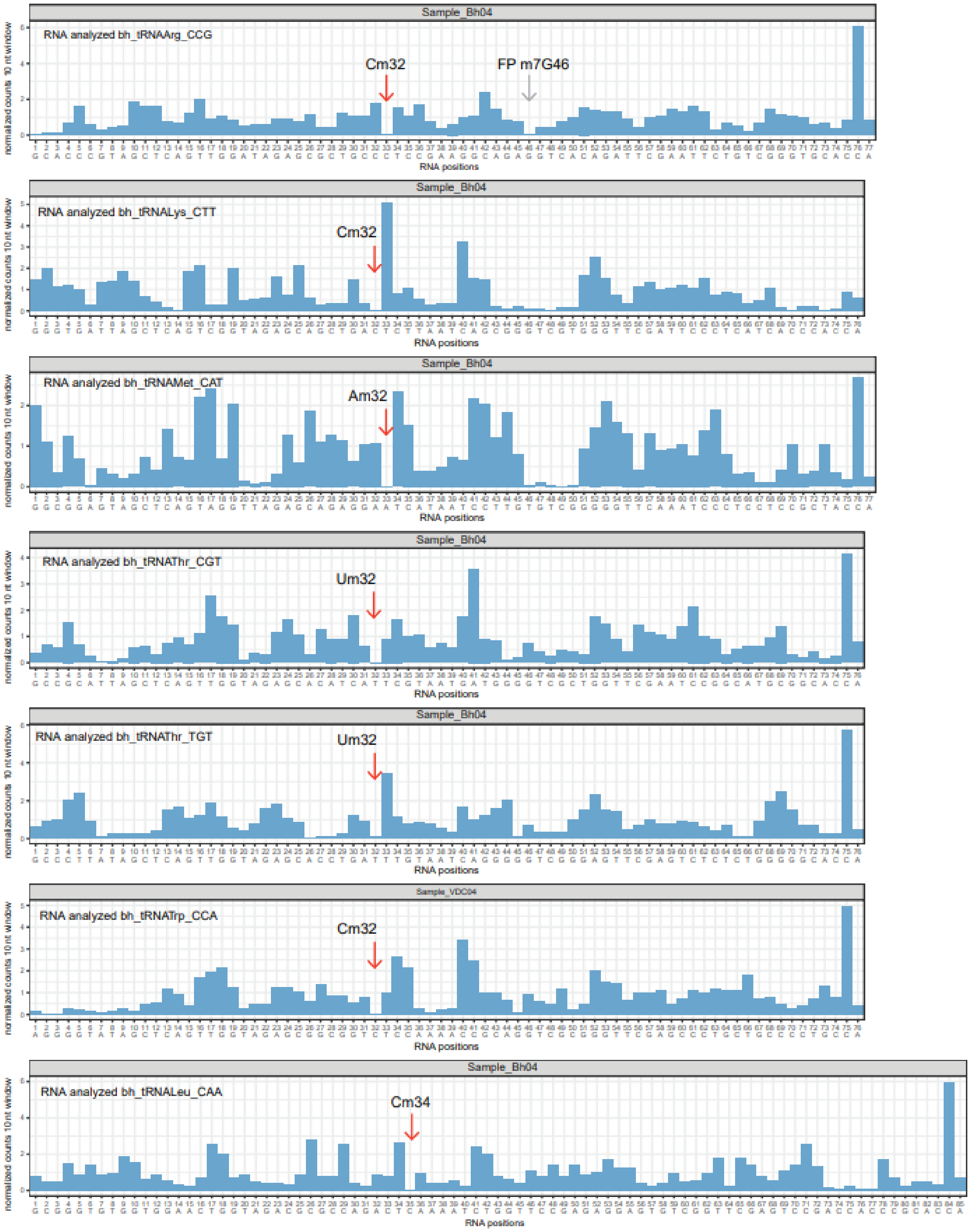


**Figure S3**. RiboMethSeq protection profiles showing alkaline hydrolysis-protected Nm residues in *B. henselae* tRNAs. Cumulative 5'- and 3'-counts normalized to the median in the 11 nt window are shown. Red arrows indicate protected Nm34, and Nm34 protection resulting from m$^7$G46 is shown by a grey arrow (tRNAArg_CCG, top trace). Sequence of tRNA species with consecutive numbering of tRNA nucleotides is shown at the bottom.

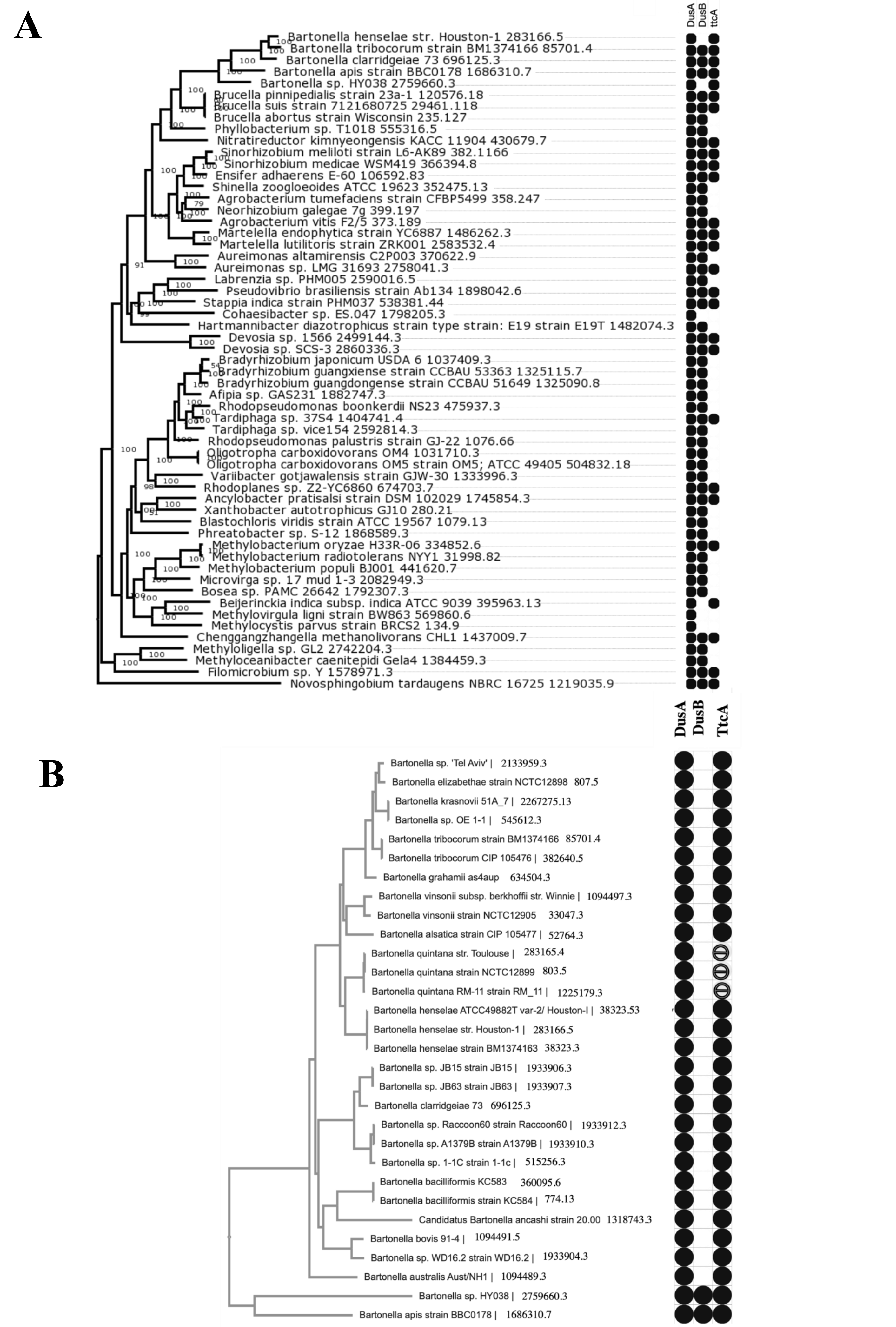


**Figure S4. DusA, DusB , and TtcA distribution in *Hyphomicrobiales* (A) and *Bartonellaceae* (B).** Gene presence and absence information for DusA, DusB, and TtcA were extracted from BV-BRC using the enzyme full names: "tRNA-dihydrouridine(20/20a) synthase," "tRNA-dihydrouridine synthase DusB," and "tRNA-(cytosine32)-2-thiocytidine synthetase TtcA," respectively. Gene presence-absence data were determined from a subset of genomes with complete sequences, and these patterns were cross-verified using NCBI Protein Blast (pBlast). BV-BRC genome IDs associated with each data point in the trees are provided for accurate identification and comparison besides the strain name and in Table S10.

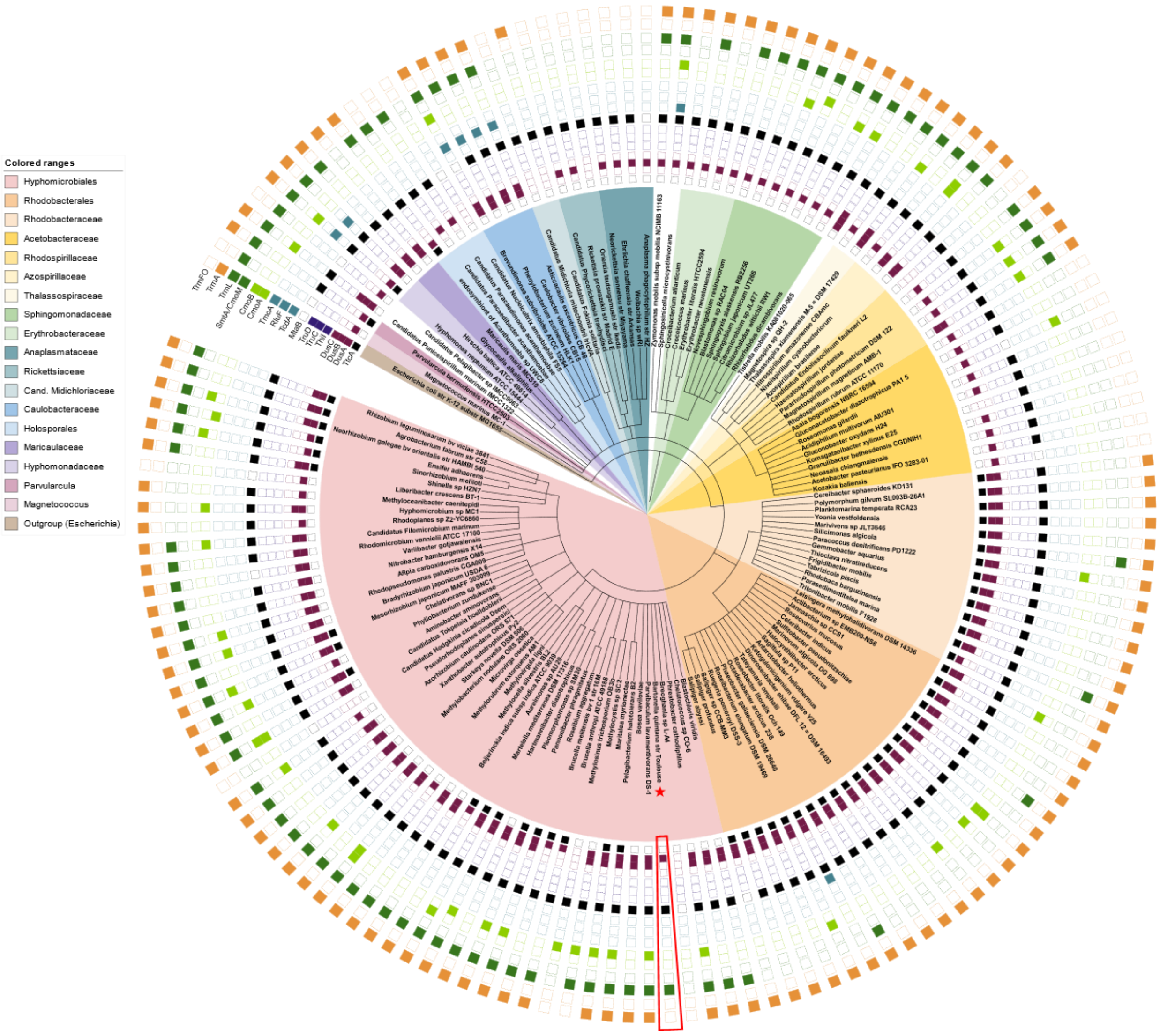


**Figure S5.** Phylogenetic tree of benchmark Alphaproteobacteria. KEGG Orthology absence-presence data is mapped to each organism for the following KOs: TtcA (K14058), DusA (K05539), DusB (K05540), DusC (K05541), ThiI (K03151), TruC (K06175), TruD (K06176), MtaB (K18707), TcdA (K22132), RluF (K06182), TmcA (K06957), CmoA (K15256), CmoB (K15257), CmoM (K06219), TrmL (K03216), TrmA (K00557), and TrmFO (K04094). Each row of mapped binary data is labeled accordingly. *Escherichia coli* K-12 MG1655 was chosen as the tree's representative outgroup. Color ranges denote subclades of interest; colors and their corresponding subclades are listed above in the range key. *Bartonella quintana* str. Toulouse is denoted in the tree by a red star, its corresponding data boxed in red. The data to generate the tree is given in Table S11, the full tree can be seen at https://itol.embl.de/tree/8216522107478961703704676
